# Leach’s Storm-Petrels fledge on the full moon and throughout the lunar cycle

**DOI:** 10.1101/2023.08.04.552000

**Authors:** Sydney M. Collins, April Hedd, William A. Montevecchi, Tori Burt, David R. Wilson, David A. Fifield

## Abstract

Many seabirds are attracted to anthropogenic light, and the risk is greater for recent fledglings. Lunar phase predicts the probability of stranding, but it remains uncertain whether lunar phase is associated with when young seabirds fledge. Fledging behaviour of nocturnal, burrowing seabirds can be difficult to monitor using traditional methods but can provide insight into environmental factors that influence the risk of stranding. We used passive integrated transponder tags to monitor the fledging dates and times of Leach’s Storm-Petrel chicks from four breeding seasons at a major colony in Newfoundland and Labrador, Canada. We also assessed whether lunar phase associated with fledging date. The median fledge time was 2.4 h after sunset (1.4 - 12.4 h). The median fledge date was 10 October, and fledging dates ranged from 13 September to 13 November. Most importantly, Leach’s Storm-Petrels chicks did fledge during the full moon. These results provide insight into why storm-petrels are less attracted to anthropogenic light during high levels of natural illumination and can be used to inform periods of higher risk for stranding, thus allowing better concentration of conservation efforts.

## INTRODUCTION

Leach’s Storm-Petrels (*Hydrobates leucorhous*) are small, nocturnal seabirds that breed in burrows in large colonies around the northern hemisphere, with most of the population in Atlantic Canada [1]. Despite their abundance (millions of birds), the Atlantic populations are declining rapidly [2–7]. Factors thought to contribute to these declines include pollution, predation, climate change, habitat modification, and attraction to anthropogenic light [2,8]. Light attraction appears to be particularly problematic for Leach’s Storm-Petrels in Atlantic Canada; thousands of birds strand annually around coastal towns, industrial buildings, and offshore oil rigs [9–12]. The vast majority of stranded storm-petrels are recent fledglings and juveniles [9–11]. Of particular concern is the episodic occurrence of mass stranding events, in which hundreds to thousands of fledgling storm-petrels strand within hours or days at a single site [11].

Although the causes of mass strandings among storm petrels are unknown, lunar phase is suspected to play a role [10]. Previous studies have observed that, on nights with a full moon, fewer birds tend to strand [9–11], and adult storm-petrels tend to be less active at the colony [13–16]. Together, these results suggest that storm-petrels avoid fledging on a full moon [17], yet to our knowledge, no study has assessed this hypothesis.

Ascertaining the factors that predict fledging of Leach’s Storm-Petrels may help predict mass stranding events, but monitoring their fledging behaviour is difficult. First, storm-petrels are nocturnal [1], so our ability to observe the time and date of fledging is limited. Second, chicks may stand outside or leave the burrow for several hours or days before returning again [1], so finding a burrow empty during a single check does not necessarily indicate that a chick has fledged.

To circumvent these challenges, we used passive integrated transponder (PIT) tags to remotely monitor fledging dates and times of Leach’s Storm-Petrel chicks. Our specific objectives were to determine (1) the peak and range of fledging date and time and (2) whether fledging is associated with lunar phase. We predicted that storm-petrel chicks do not fledge when the moon is fuller [17]. Knowledge of fledging time and any coordination with environmental factors will enhance our ability to predict mass-stranding events, thus allowing more concentrated monitoring during the periods of highest risk.

## Methods

### Field Methods

#### Field Site

We studied Leach’s Storm-Petrel chicks on Gull Island (47.26265, -52.77187), Witless Bay, Newfoundland and Labrador, Canada from 2017 to 2022. Gull Island supports approximately 180000 breeding pairs of Leach’s Storm-Petrels [18]. Chicks were monitored across 6 plots distributed along the southwestern side of the island (Figure S1).

#### PIT Tag Setup

Cylindrical glass 150 kHz PIT tags were set inside a custom 3D printed leg band (either 12 mm x 2.12 mm CoreRFID model SOK027, 0.25 g total weight, or 10 mm x 2.12 mm Cyntag model 601205-248, 0.15 g total weight) and mounted on the leg of Leach’s Storm-Petrel chicks (Figure S2 A). Each chick was banded with a unique stainless steel identification band on the other leg, weighed, and measured for wing chord length. Chick banding began in late August or early September of each year. Leach’s Storm-Petrels have high hatching asynchrony [1], so not all chicks that were banded were large enough to be equipped with a PIT tag. These chicks were revisited later in the season when possible or were not included in this study. Tag reader antennae (Figure S2 B) consisted of wire coils wrapped around custom 3D-printed plastic cylinders (72 mm diameter by 20 mm deep) and a tuning capacitor. The antennae were inserted into the mouth of the burrow and secured in the ground using garden stakes. Each antenna was connected to a custom-made circuit board housed inside a Pelican Case, which recorded the time and identification of the bird as it passed through the antenna. Video footage indicates that the antennae did not impede the storm-petrels’ movement in or out of the burrow. The circuit board recorded system information (e.g., antenna frequency, battery voltage, etc.) and re-tuned the antenna every 30 minutes, which allowed us to identify the occurrence of system failures.

### Verification of Fledging

In most cases, the final read for each chick was considered the time and date at which the chick fledged. We could not physically verify fledging because (1) dead chicks sometimes become buried in the burrow chamber and cannot be felt by researchers during burrow inspections (pers. observation), (2) chicks may die or be depredated outside the burrow while exploring [1], and (3) researcher access to the colony can be limited during the fledging period due to inclement weather. We, therefore, estimated the age of each chick at banding to determine whether the chick was old enough to fledge by the date of the last read. We estimated chick age from wing length using an equation derived by R.A. Mauck (unpubl.). A chick was assumed to have fledged if its estimated age at last read exceeded 56 days, as this represents the minimum fledging age observed across multiple colonies [1].

### Statistical Methods

All analyses were conducted using R version 4.2.2 [19]. Dates were converted to day of year using the “yday” function of the package *lubridate* [20], *and times were calculated as hours after sunset using “as*.*ITime” and “daylength” from data*.*table* [21] *and chillR* [22], respectively. Summary statistics were calculated for the fledging dates and times. ANOVAs (or Kruskall-Wallace tests when data were non-normal) were used to determine differences among years in mean fledging date and time. Lunar phase [23] was categorized into quarters. A Chi-squared goodness of fit test was used to determine if the proportion of birds that fledged during each lunar phase differed from the expected proportions (i.e., the proportion of days within the fledging period of each illumination condition). Similar analyses examining the interactive effects of cloud cover [24,25] and lunar phase were conducted. These analyses are included in the supplementary material due to low statistical power and low-resolution cloud cover data (Figures S3 and S4).

## RESULTS

From 2017 to 2022, 122 chicks were tracked using PIT tag technology (Table 1). Based on the calculation of chick age at fledging (R.A. Mauck unpubl.) and a minimum fledge age of 56 days [1], 2 chicks were deemed too young to fledge at the time of their final read and were thus eliminated from the sample (n = 120). The median fledge date of all chicks was 10 October (IQR: 15.3 days, range: 13 September - 13 November), and the median fledge time was 2.4 h after sunset (IQR: 1.3, range: 1.4 - 12.4 h) (Table 1, Figure 1, Figure S5).

**Table 1.** Summary statistics of the fledge date and time ± IQR of Leach’s Storm-Petrel chicks from Gull Island, Witless Bay, Newfoundland and Labrador, Canada. Ranges are reported in brackets.

| Year | Sample Size | Median Time (hours) | Median Time Past Sunset (hours) | Median Date (days) |
| --- | --- | --- | --- | --- |
| 2017 | 30 | 22:00 ± 1.3 h<br>(19:17 - 05:30) | 2.2 ± 1.2 h<br>(1.5 - 12.4) | 11 Oct ± 14.3 days<br>(19 Sept - 29 Oct) |
| 2018 | 41 | 22:15 ± 0.9 h<br>(18:47 - 03:57) | 2.4 ± 0.7 h<br>(1.7 - 10.1) | 8 Oct ± 17.0 days<br>(19 Sept - 31 Oct) |
| 2021 | 9 | 22:20 ± 1.1 h<br>(19:11 - 01:11) | 2.4 ± 1.3 h<br>(1.8 - 7.6) | 29 Sept ± 13.0 days<br>(25 Sept - 18 Oct) |
| 2022 | 40 | 22:12 ± 1.9 h<br>(18:54 - 05:00) | 2.6 ± 1.9 h<br>(1.4 - 11.2) | 11 Oct ± 18.8 days<br>(13 Sept - 13 Nov) |
| All | 120 | 22:11 ± 1.3 h<br>(18:47 - 05:30) | 2.4 ± 1.3 h<br>(1.4 - 12.4) | 10 Oct ± 15.3 days<br>(13 Sept - 13 Nov) |

**Figure 1.**
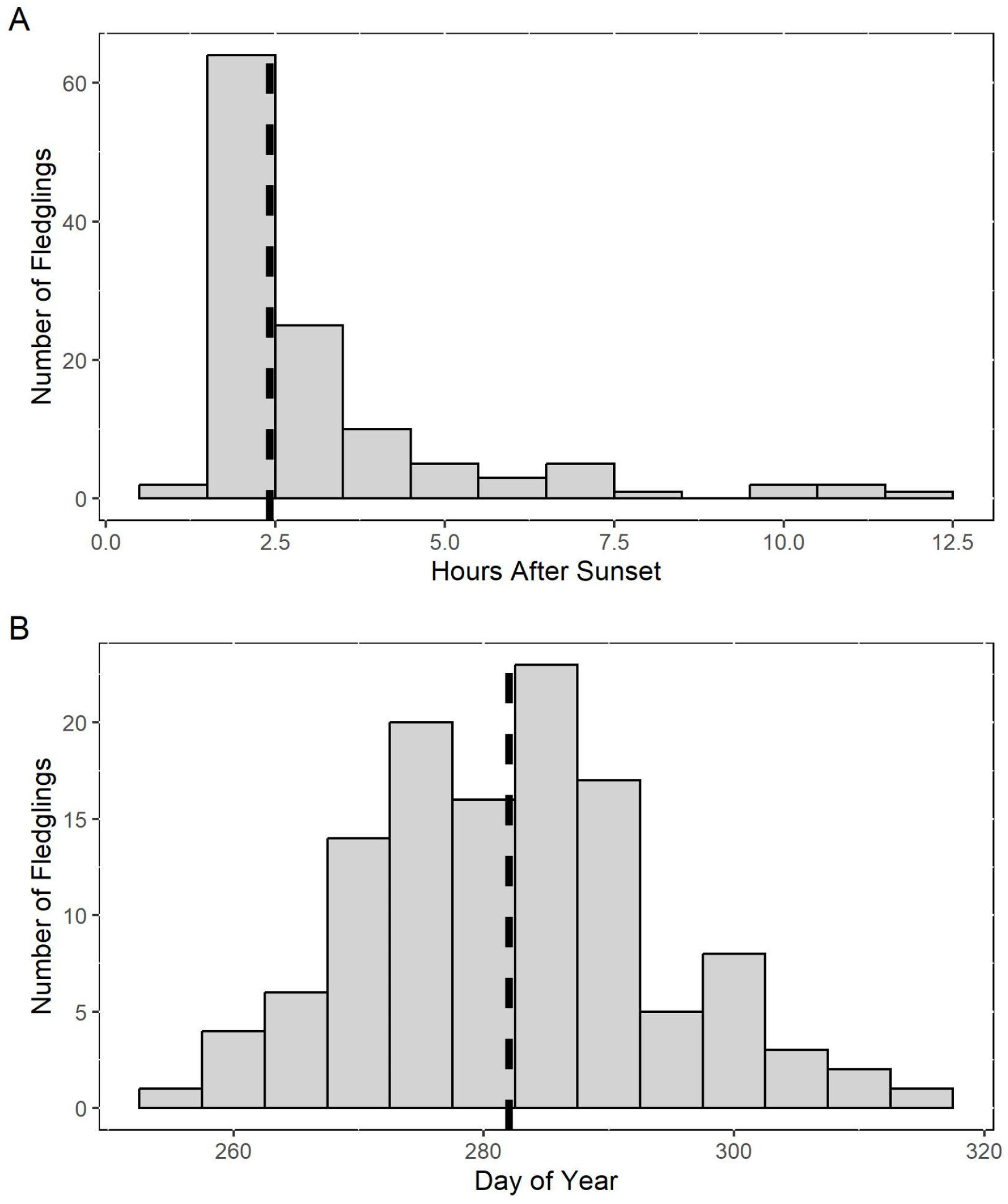
Histograms of the A) time after sunset and B) day of year that Leach’s Storm-Petrel chicks fledged from Gull Island, Witless Bay, Newfoundland and Labrador, Canada. The dashed black line is the median.

There were no significant differences in the time past sunset (Kruskall-Wallace χ^2^ = 1.42, df = 3, p = 0.70) or date (F = 2.62, df = 3, p = 0.054) of fledging among years. The chisquared test for goodness of fit found that the number of chicks that fledged during each lunar phase differed significantly from the expected number of fledglings in each condition (χ^2^ = 11.63, df = 3, p = 0.0087; Figure 2; Figure S6). One-proportion z-tests revealed that fewer than expected birds fledged during the second quarter, and more than expected birds fledged during the third quarter (Table 2).

**Table 2.** Results of the one-sample proportion tests examining the expected and observed number of Leach’s Storm-Petrel chicks to fledge under different lunar phases.

| Lunar Phase | Expected number of Fledglings | Observed Number of Fledglings | Chi-squared Value | P-value |
| --- | --- | --- | --- | --- |
| First Quarter | 36 | 34 | 0.15 | 0.70 |
| Second Quarter | 20 | 10 | 5.12 | 0.023 |
| Third Quarter | 20 | 32 | 7.4 | 0.0065 |
| Fourth Quarter | 44 | 44 | 0 | 1 |

**Figure 2.**
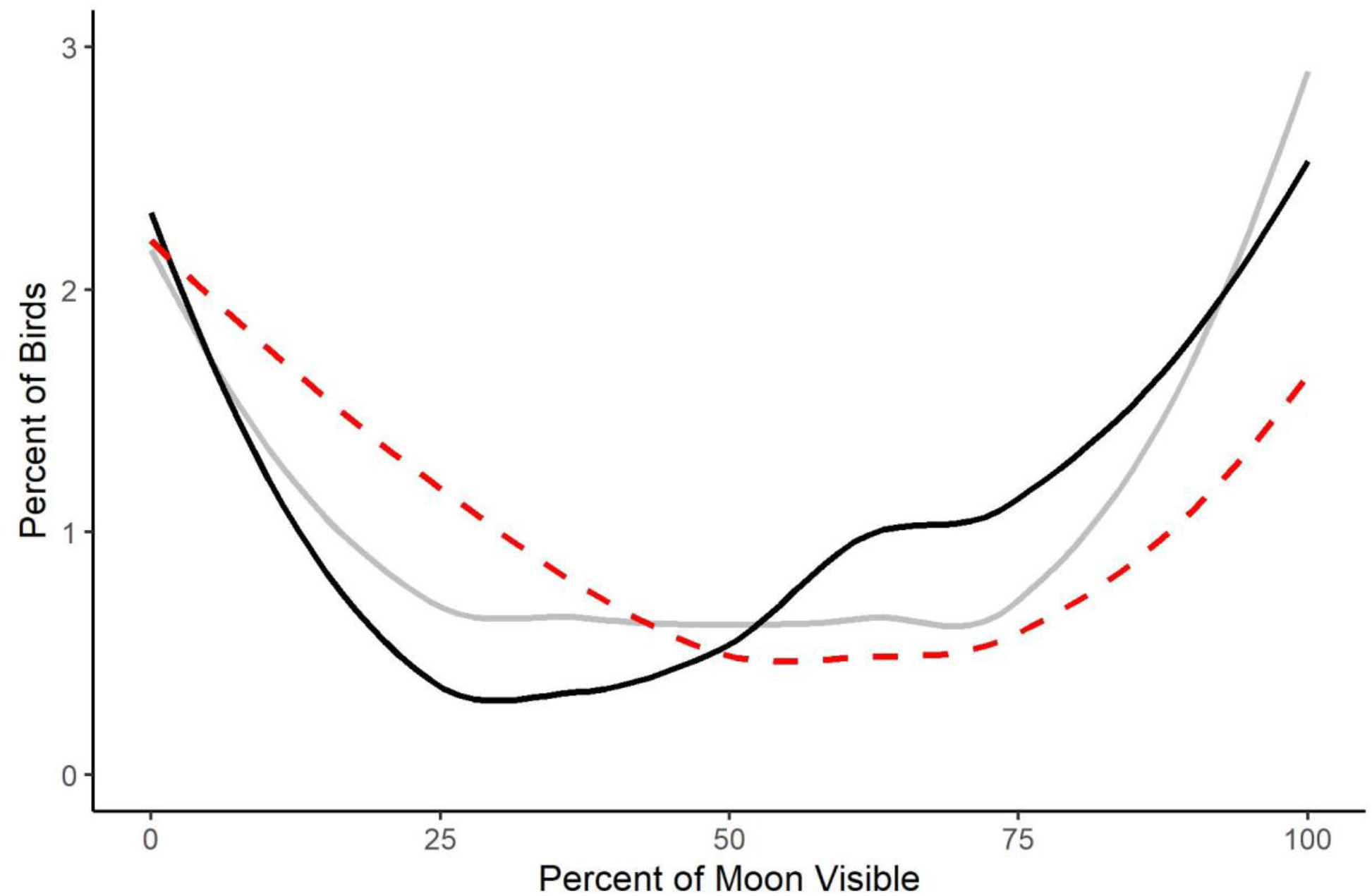
Line plot of the observed percentage of fledglings (solid black line) and stranded birds (red dotted line) occurring during different lunar phases. These are compared with the expected proportion of fledglings or stranded birds (grey solid line) should birds fledge/strand randomly relative to lunar phase. Strandings data ([11], T.V. Burt unpubl. data) were collected from 2019 to 2022 in Bay de Verde, Newfoundland and Labrador, Canada. Mass stranding events (>100 birds stranded in one night) were excluded from this plot.

## DISCUSSION

Using data from passive integrated transponder technology, we determined the median fledging date and time of Leach’s Storm-Petrel chicks to be 10 October 2.4 h after sunset. Fledging dates ranged from mid-September to mid-November, which aligns with previous reports from colonies in Atlantic Canada [1]. These dates also align with periods of peak strandings reported for Leach’s Storm-Petrels in Newfoundland [9–12]. Studies documenting stranded storm-petrels report that the majority of birds which strand during this period are fledglings [10,11], so it is very likely that the recovered birds are recent fledglings from nearby colonies.

We observed fledging times very close to sunset in this study (Figure 1, Table 1), which is interesting considering peak activity at the colony of adult Leach’s Storm-Petrels is much later in the night [26]. It is unknown for how long storm-petrel fledglings remain at the colony after departing their burrow. These early fledging times, however, concur with findings from surveys of stranded fledgling Procellariiformes (including Leach’s Storm-Petrels), which observed peak stranding within a few hours of sunset [27,28].

Contrary to our hypothesis, storm-petrels fledged during the full moon (Figure 2). This result is surprising for two reasons. Adults tend to reduce their activity at the colony during the full moon [13–16], which may be a strategy to avoid predation. As well, fewer Leach’s Storm-Petrels tend to strand during the full moon [9,10]. Storm-petrel chicks fledging during the full moon may indicate that attraction to anthropogenic light is tempered by moon illumination.

Several hypotheses seek to explain why storm-petrels and other seabirds are attracted to anthropogenic light. First, storm-petrels may orient toward anthropogenic light because they mistake it for their bioluminescent prey [29]. Second, storm-petrels may navigate using light from the moon and stars [30], so anthropogenic light may be disorienting and cause them to orient towards it. Finally, juveniles may be more susceptible to light attraction due to their underdeveloped retinas [31,32]. Storm-petrels fledging during the full moon has interesting implications for the latter two hypotheses. If storm-petrels use the light from the moon and stars to navigate, fewer storm-petrel fledglings may strand during the full moon because they can navigate better with more natural light. In addition, if fledglings and juveniles are particularly vulnerable to light attraction, greater lunar illumination will reduce the relative intensity of anthropogenic light, presumably reducing their attraction [15,27].

Storm-petrels are less active on the colony during a full moon, possibly as a method of predation avoidance [13]. At the colony, the dominant predators of Leach’s Storm-Petrels are often diurnal Charadriiformes such as Herring Gulls *(Larus argentatus*) and Great Skuas (*Stercorarius skua*) [18,33,34]. Though these predators can forage at night, they likely benefit from well-lit conditions provided by greater nocturnal illumination [13,35–37]. In response, storm-petrels may remain inside the burrow to avoid being easily detected and predated, resulting in low colony activity outside the burrows. This behaviour may be innate as other seabirds have been shown to adjust their activity based on moon illumination even in the absence of predation pressure on the colony [38,39]. Leach’s Storm-Petrel fledglings, however, depart from their burrow during a full moon, so moonlight avoidance behaviour may in fact be learned rather than instinctual. The lack of moonlight avoidance while fledging from Gull Island may be due to the fact that most gulls are no longer present at the colony when storm-petrel chicks begin to fledge [40,41], so there is no antipredator benefit to chicks to avoid fledging under a full moon. It is unknown, however, how long after fledglings depart from their burrow that they leave the colony. It is possible that fledglings remain at the colony longer during a full moon, which could explain why fewer fledglings strand at this time. Future research should track the location of fledglings during their inaugural flight to investigate the timing and conditions of departure, and to determine whether the direction of travel (towards anthropogenically lit areas or out to sea) is influenced by nocturnal illumination (i.e. [28]).

From a conservation perspective, these results indicate that rescue programs should concentrate their efforts early in the night from mid-September to mid-November, but monitoring decisions should not be based on lunar phase. Other factors like wind speed, wind direction, fog, and the brightness and colour of anthropogenic light may influence the likelihood of birds stranding [42]. Because storm-petrel fledglings do not appear to base their decision to leave the burrow on lunar phase, mass stranding events are possible during the full moon. Long-term studies of mass-stranding events, such as those conducted in Bay de Verde and Witless Bay [10,11,43], are needed to determine factors influencing the probability of mass-stranding events. This information could be used to reduce industrial lighting and flaring during high-risk conditions and periods (e.g., foggy conditions during mid-September through November) [44]. Understanding such factors will allow conservation groups to mitigate and respond to stranding events more effectively.

## Acknowledgements

We thank Krista Baker, Katie Birchard, Megan Boucher, Taylor Brown, Chantelle Burke, Greg Campbell, Lily Colston-Nepali, Joshua Cunningham, Kyle d’Entremont, Parker Doiron, Mohammad Fahmy, Michelle Fitzsimmons, Carina Gjerdrum, Bronwyn Harkness, Emma Lachance Linklater, Keith Lewis, Shannon Mawhinney, Gretchen McPhail, Marina Montevecchi, Isabeau Pratte, Cerren Richards, Greg Robertson, Pierre Ryan, Sydney Shepherd, Manon Sorais, Katherine Studholme, Michelle Valiant, Christopher Ward, Sabina Wilhelm, and Jeannine Winkel for help with fieldwork.

We further thank Mohammad Fahmy, Madeline Scevoir, and Sophie Burke for assisting with data analysis. Thank you to Rod Byrne for designing the burrow monitor circuit boards and creating the system software. Funding for this project was provided by the Natural Sciences and Engineering Research Council of Canada [2018-06872 to WAM, RGPIN-2015-03769 to DRW, and PGSD3-547858-2020 to SMC], Environment and Climate Change Canada [GCXE22C307 to WAM and internal funding], and Memorial University of Newfoundland and Labrador.

## Ethics Statement

Safe capture, handling, and banding of animals was performed under scientific permit number SC2674 and banding permit numbers 10332 U and 10559 N. Access to the Witless Bay Ecological Reserve was permitted by the Natural Areas Program, Newfoundland and Labrador Department of Environment and Climate Change.

