## Supplementary Material for "Leach’s Storm-Petrels fledge on the full moon and throughout the lunar cycle"

**Appendix**

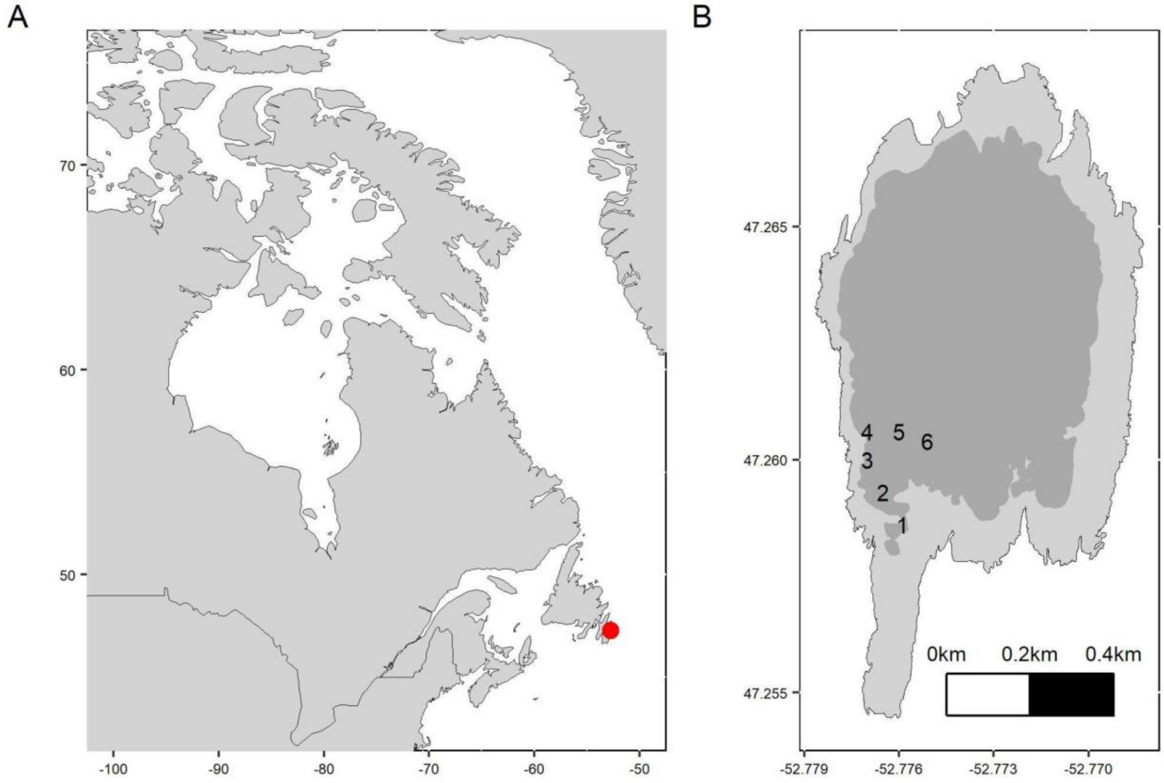


Figure S1. Maps indicating (A) the location of Gull Island relative to Eastern Canada and (B) the location and habitat of the six PIT tag plots on Gull Island, where dark grey represents forest and light grey represents grassy slopes.

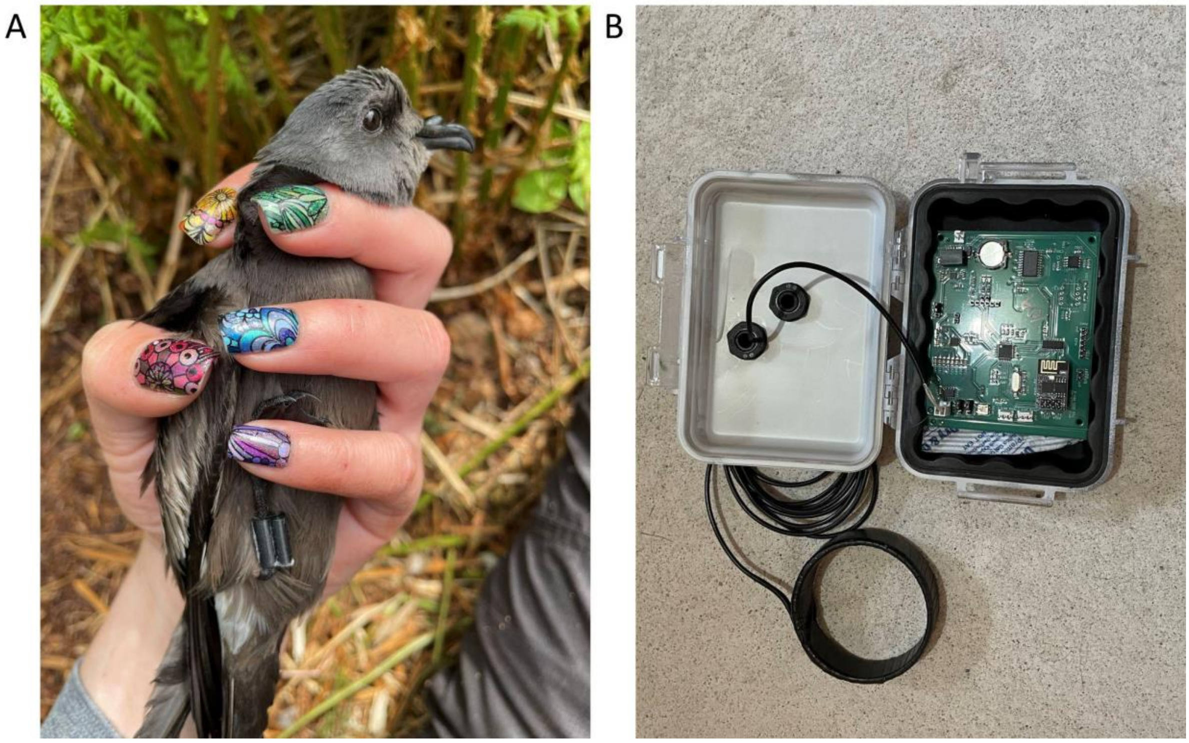


Figure S2. (A) Leach's Storm-Petrel equipped with a PIT tag contained within a custom 3D-printed leg band. (B) PIT tag system including the tag reader (black circle) which is inserted into the mouth of the burrow.

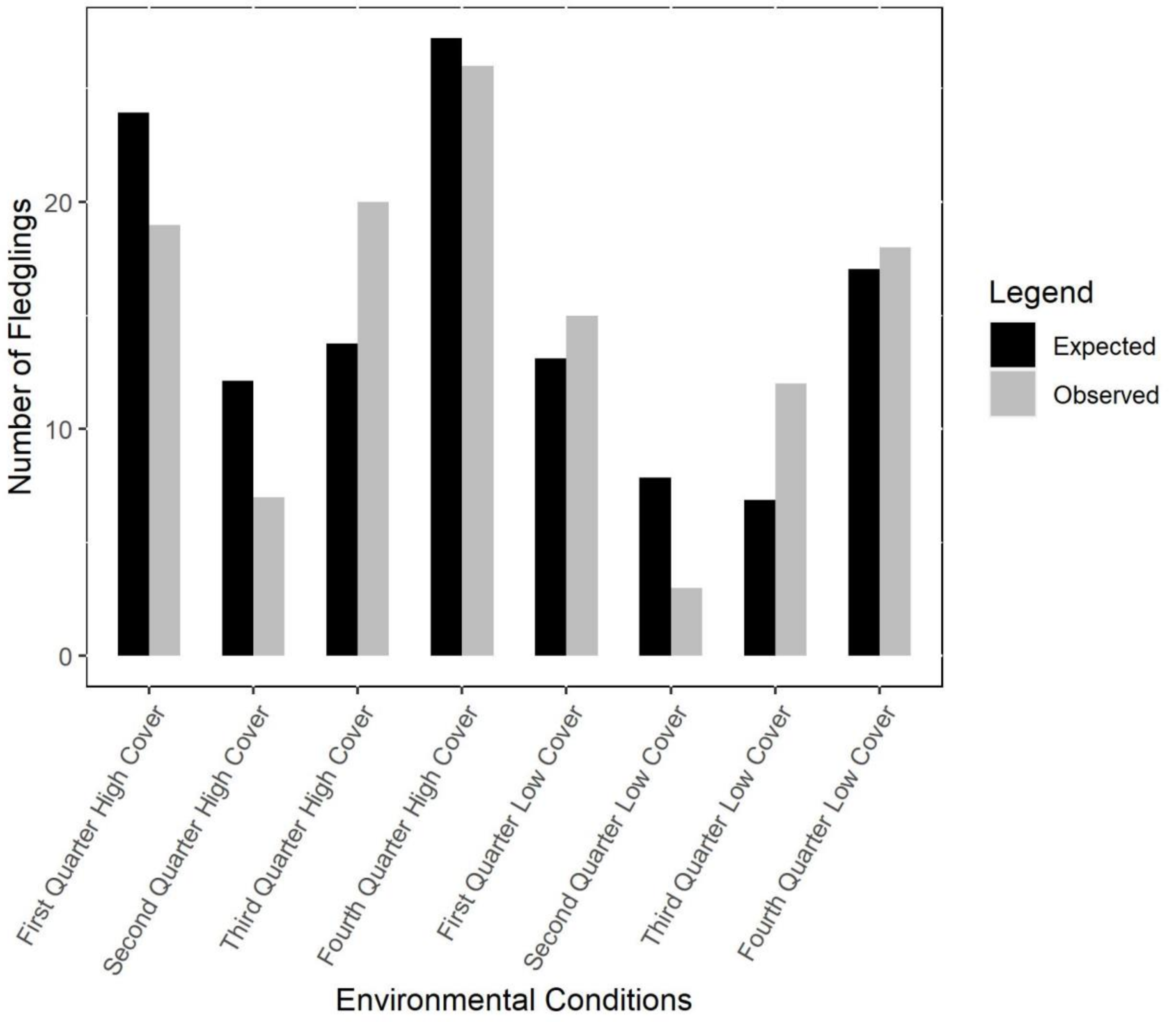


Figure S3. Observed vs. expected number of flegdlings given the moon illumination and cloud cover ($\chi2 = 13.38$, df = 7, $p = 0.063$).

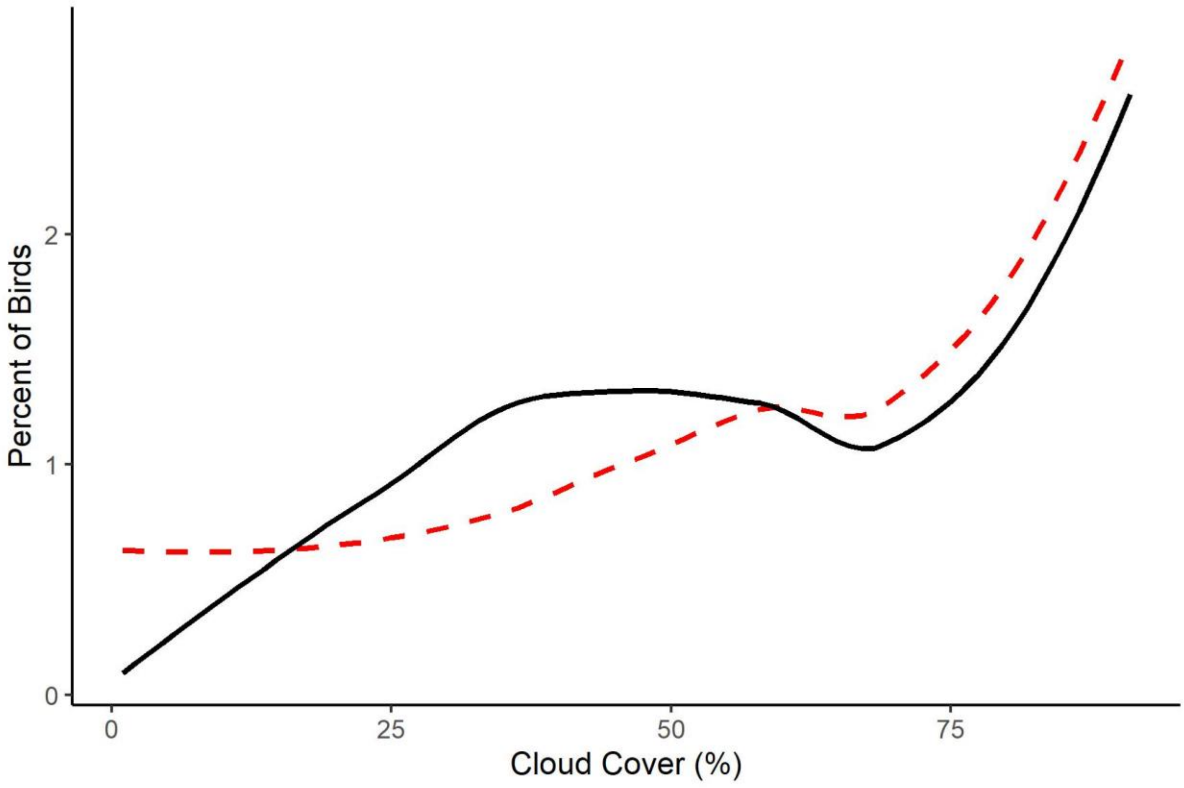


Figure S4. Observed (black solid line) versus expected (red dashed line) proportion of fledglings given cloud cover ($\chi^2$ = 4.43, df = 3, p = 0.22).

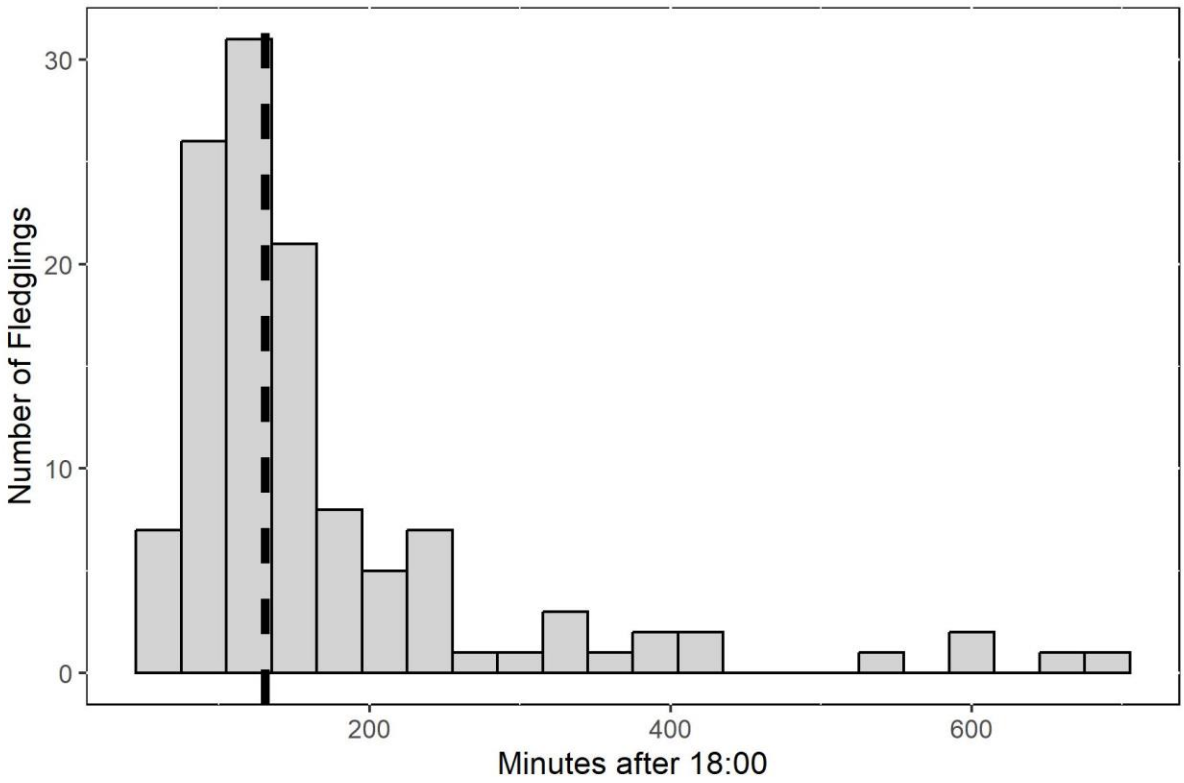


Figure S5. Histogram of the time of day (minutes after 18:00) that Leach's Storm-Petrel chicks fledged. The black dashed line is the median (22:11).

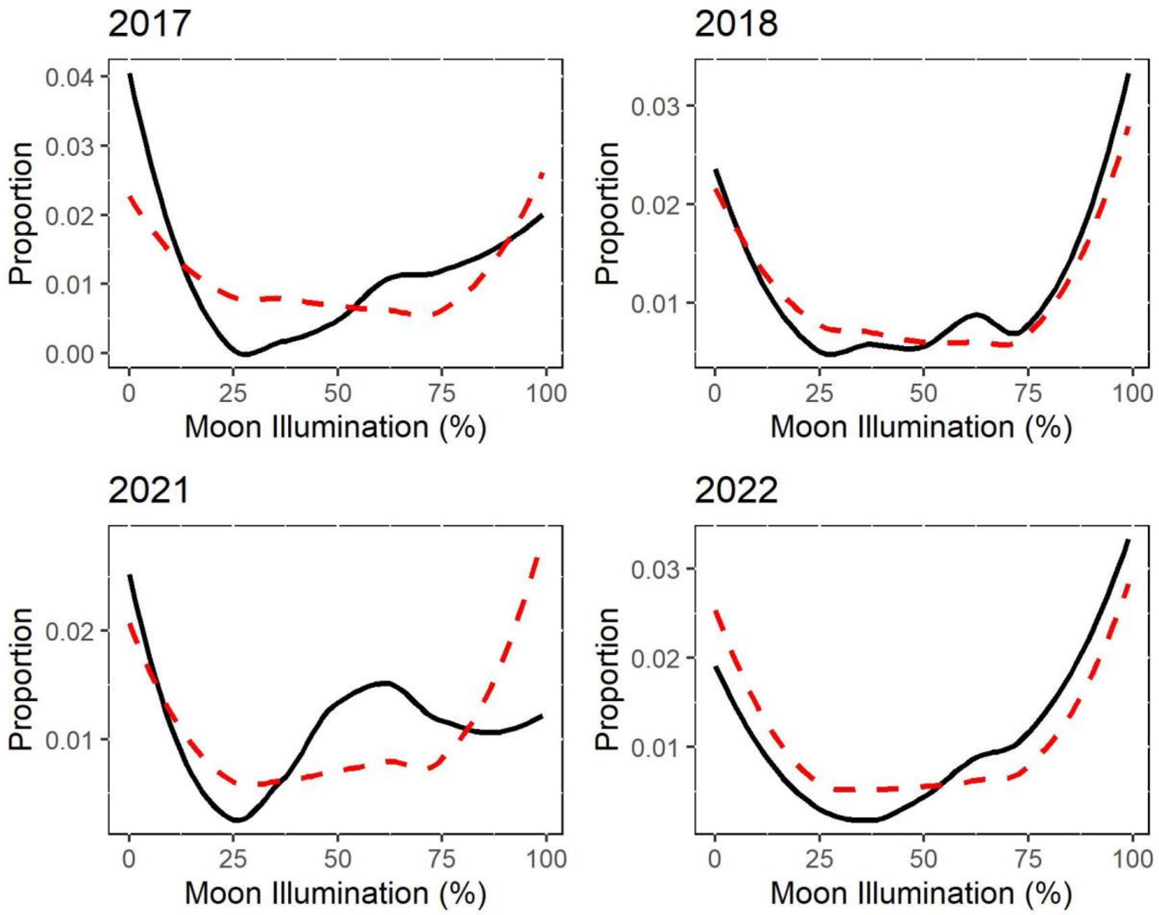


Figure S6. Observed (black solid line) versus expected (red dashed line) proportion of fledglings given moon illumination for each year of the study.
